# Multidimensional hydrogel models reveal endothelial network angiocrine signals increase glioblastoma cell number, invasion, and temozolomide resistance

**DOI:** 10.1101/2020.01.18.911396

**Authors:** Mai T. Ngo, Elijah Karvelis, Brendan A.C. Harley

## Abstract

Glioblastoma is the most common primary malignant brain tumor. The tissue microenvironment adjacent to vasculature, termed the perivascular niche, has been implicated in promoting biological processes involved in glioblastoma progression such as invasion, proliferation, and therapeutic resistance. However, the exact nature of the cues that support tumor cell aggression in this niche are largely unknown. Soluble angiocrine factors secreted by tumor-associated vasculature have been shown to support such behaviors in other cancer types. Here, we exploit macroscopic and microfluidic gelatin hydrogel platforms to profile angiocrine factors secreted by self-assembled endothelial networks and evaluate their relevance to glioblastoma biology. Aggregate angiocrine factors support increases in U87-MG cell number, migration, and therapeutic resistance to temozolomide. We also identify a novel role for TIMP1 in facilitating glioblastoma tumor cell migration. Overall, this work highlights the use of multidimensional hydrogel models to evaluate the role of angiocrine signals in glioblastoma progression.

**Insight, Innovation, and Integration:** Glioblastoma progression is linked to interactions between tumor and vascular cells, which can influence invasion and therapeutic response. In co-culture studies to investigate tumor-vascular crosstalk, endothelial cells often are not presented in three-dimensional structures mimicking vasculature and the exact identity of secreted factors is not explored. Here, we use tissue engineering strategies to generate three-dimensional endothelial networks from which to collect soluble angiocrine signals and assess the impact of these signals on glioblastoma behavior. Furthermore, we use secretomic analysis to identify specific factors influencing glioblastoma invasion. We identify a novel role for TIMP1 in supporting glioblastoma migration and demonstrate that soluble angiocrine signals support chemoresistance to temozolomide.

## 1. Introduction

Glioblastoma (GBM) is the most common primary malignant brain tumor^1^. With a median survival of 15 months and a five-year survival rate of 5%^2^, poor patient survival is linked to rapid and invasive spreading of GBM cells throughout the brain. There is an urgent need to develop an improved understanding of the mechanisms governing GBM invasion, proliferation, and drug resistance in order to identify novel therapeutic targets. The physical nature and biochemical composition of the extracellular matrix (ECM), as well as crosstalk between GBM tumor cells and surrounding vascular, immune, and stromal cells contribute to tumor cell growth, migration, proliferation, and therapeutic response^3,4^.

GBM tumor cells have been observed to reside in perivascular niches (PVN), which are tissue microenvironments surrounding vasculature, within the brain^5,6^. GBM cells migrate toward and associate closely with blood vessels as they spread throughout the brain in an act called co-option^7,8^. Co-option ultimately results in damage to the blood vessels, which then regress^9,10^, triggering formation of hypoxic zones which initiate pro-angiogenic signaling to promote new capillary formation at the tumor periphery^11^. Tumor cells will then migrate towards the newly-formed vasculature. In this manner, vasculature is believed to contribute significantly to GBM progression by facilitating migration and by modulating nutrient and oxygen supply. Insight into the cues that initiate vessel co-option will be beneficial for determining strategies to mitigate tumor invasion. Furthermore, recent studies in regenerative, stem cell, and cancer biology suggest that vasculature can instruct the behavior of surrounding cells via angiocrine signaling cues derived from the cells that make up vasculature, namely endothelial cells and supporting mural cells (e.g. pericytes, smooth muscle cells)^12-15^. Understanding how signaling cues from vasculature affect tumor cell behavior and the underlying mechanisms that govern the crosstalk between vasculature and tumor cells can provide insights into new therapeutic strategies, particularly because the PVN has been associated with the maintenance of stem-like and therapeutically-resistant GBM cells that facilitate disease recurrence^6,16-18^.

In vitro co-culture, Transwell, or conditioned media experiments have been used to investigate the crosstalk between endothelial and GBM cells^19-21^. However, these studies are often hindered by the inability of the endothelial cells to assemble into structures resembling vasculature. Furthermore, these experiments are often performed in 2D, which has limited relevance to native 3D tissue. Biomaterial platforms can be used to overcome these limitations. Specifically, endothelial and mural cells (e.g. fibroblasts, mesenchymal stem cells) can be encapsulated within biomaterials to generate three-dimensional vascular networks^22-24^. Engineered vasculature has been shown to deposit basement membrane proteins and express tight junctions, and mural cells assume perivascular positions in these cultures similar to native pericytes^25-26^. These characteristics suggest that biomaterials can be used to develop physiologically-relevant vascular networks. Biomaterials have also been used to generate 3D culture models for tumors, with numerous studies demonstrating that biomaterial cultures better recapitulate tumor invasion, proliferation, and therapeutic response compared to two-dimensional culture^27-30^.

We have previously described a gelatin hydrogel platform for co-culturing endothelial cells, fibroblasts, and GBM tumor cells to model the GBM PVN^31^. Here, we specifically investigate the role of the secretome of 3D self-assembled endothelial networks in influencing the behavior of U87-MG tumor cells encapsulated in gelatin hydrogels. We collect and profile the identities of proteins secreted by networks formed from endothelial cells and fibroblasts cultured in gelatin hydrogels^32^. We subsequently determine the effects of the secretome on U87-MG cell number, metabolic activity, migration, and response to temozolomide (TMZ), the standard chemotherapy for GBM.

## 2. Materials and Methods

### 2.1. Cell Culture

U87-MG cells were purchased from ATCC (Old Town Manassas, VA) and passaged less than 10 times. Dulbecco’s Modified Eagle Medium (DMEM) with 10% fetal bovine serum (FBS) and 1% penicillin/streptomycin was used to culture U87-MG cells. Human umbilical vein endothelial cells (HUVECs) and normal human lung fibroblasts (NHLFs) were purchased from Lonza (Walkersville, MD) and used before passage 6 and passage 8 respectively. HUVECs were cultured in Endothelial Growth Medium 2 (EGM-2) and NHLFs were cultured in Fibroblast Growth Medium 2 (FGM-2) (Lonza, Walkersville, MD). All cells were cultured in incubators at 37 °C and 5% CO_2_. Cells were tested for mycoplasma using the MycoAlert Mycoplasma Detection Kit (Lonza, Walkersville, MD), and plasmocin (0.2% v/v) (Invivogen, San Diego, CA) was added to media to prevent mycoplasma contamination.

### 2.2. Conditioned Media Collection from Perivascular Hydrogel Cultures

Methacrylamide-functionalized gelatin (GelMA) was synthesized as previously described, and degree of functionalization (DOF) was determined using ^1^HNMR^33^. GelMA with a DOF of ∼60% was used for this study. Hydrogels containing endothelial networks were formed by dissolving GelMA (5 wt%) in PBS at 65 °C. Lithium acylphosphinate (LAP) was subsequently added (0.1% w/v) as a photoinitiator. A mixture of HUVECs and NHLFs (2:1 NHLF:HUVEC, 2 × 10^6^ NHLF/mL) was re-suspended in the GelMA solution, and the resulting cell suspension was pipetted into Teflon molds (5 mm diameter, 1 mm thick). Hydrogels were formed after photopolymerization for 30 s using a UV lamp (λ = 365 nm, 5.69 mW/cm^2^). Hydrogels were deposited into 48-well plates and cultured for six days using EGM-2 media, with daily media changes. After six days, the media was switched to Endothelial Basal Medium 2 (EBM-2) (Lonza, Walkersville, MD) with 2% FBS. EBM-2 is EGM-2 without the added SingleQuot supplements. Conditioned media was collected after 24 hours, filtered using a 0.2 µm syringe filter, and stored at −20 °C until further use (**Figure 1**).

**Figure 1.**
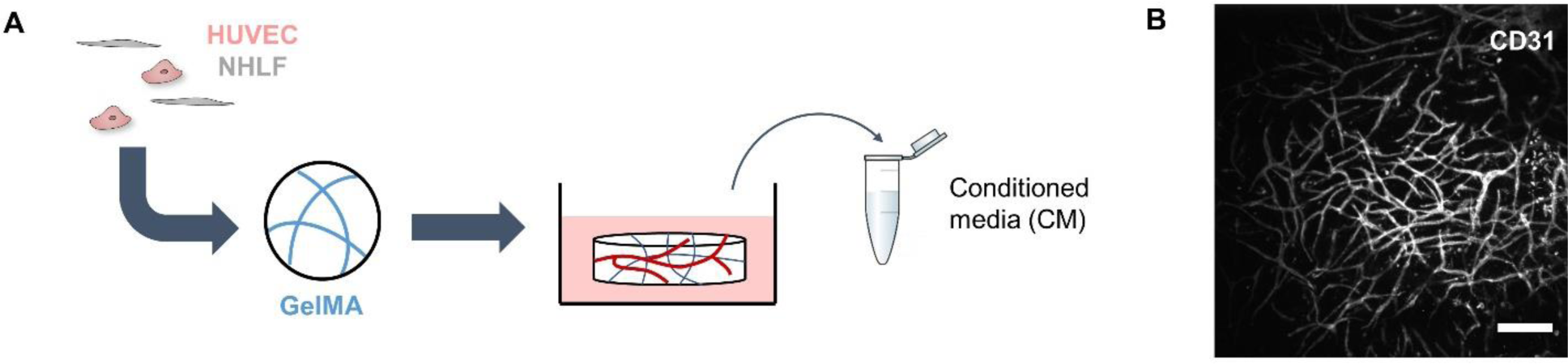
Schematic detailing the collection of conditioned media. A) Endothelial networks are formed in GelMA hydrogels by co-culturing HUVECs and NHLFs for one week. B) Endothelial networks formed after a week in culture can be visualized by staining for CD31. Scale bar = 200 µm.

### 2.3 Cytokine Array

The composition of the conditioned media was profiled using the Proteome Profiler Angiogenesis Array Kit (R&D Systems, Minneapolis, MN). 0.5 mL of media was used per assay. Membranes were imaged using an ImageQuant LAS 4010 (GE Healthcare Bio-Sciences, Pittsburgh, PA). Intensities of the cytokine spots were quantified using ImageJ (NIH, Bethesda, MD) and normalized to the intensity of the positive reference spots. Proteins with cytokine spots with intensities greater than 10% of the average intensity of the positive reference spots were identified as present in the conditioned media. Conditioned media samples from three independent experiments were analyzed. GlioVis (http://gliovis.bioinfo.cnio.es/) was used to compare expression levels of genes encoding for the identified proteins between GBM specimens and normal brain specimens collected by The Cancer Genome Atlas^34^. An ELISA (Cat# DTM100, R&D Systems, Minneapolis, MN) was used to confirm the presence and concentration of TIMP1.

### 2.4 Growth and Metabolic Activity Assays

U87-MG cells (2 × 10^6^/mL) were cultured in 5 wt% GelMA hydrogels crosslinked by photopolymerization (λ = 365 nm, 5.69 mW/cm^2^, 30 s exposure time) with 0.1% w/v LAP. Hydrogels were cultured for up to seven days in 1:1 DMEM + 2% FBS: EBM-2 + 2% FBS (*EBM2*) or 1:1 DMEM + 2% FBS: Conditioned Media (*CM*), with daily media changes. To measure changes in cell number, hydrogels were degraded using papain (Sigma Aldrich, St. Louis, MO) for 24 hours, and the lysates were mixed with Hoescht 33258 dye (Thermo Fisher Scientific, Waltham, MA). Fluorescence was read using a Tecan Infinite M200 Pro microplate reader (Switzerland). Changes in metabolic activity were assessed using an MTT assay (Thermo Fisher Scientific, Waltham, MA). Hydrogels were each incubated with 20 µL MTT and 200 µL DMEM + 2% FBS for four hours, after which the solutions were replaced with DMSO and the hydrogels were further incubated for 18 hours. Absorbance was assessed using a Biotek Synergy HT microplate reader (Winooski, VT). Proliferation and metabolic activity for each sample were first expressed as fold change compared to Day 0. Then, fold change values for *CM* samples were normalized to the average fold change value for *EBM2* samples within each independent experiment to account for varying baseline readings.

### 2.5 Invasion Assay

U87-MG migration in response to conditioned media or individual proteins was assessed using a microfluidic invasion assay. U87-MG cells were incubated in 10 µM Cell Tracker Green CMFDA (Thermo Fisher Scientific, Waltham, MA) for 30 minutes. 5 wt% GelMA was photopolymerized in the central gel channel of AIM Biotech 3D Cell Culture chips (Singapore) using UV light (λ = 365 nm, 8.39 mW/cm^2^, 25 s exposure time). After hydrating the side media channels with DMEM + 2% FBS for one hour, 4 × 10^4^ U87-MG tumor cells suspended in 15 µL of DMEM + 2% FBS were injected into one media channel, and all media ports were subsequently filled with 50 µL of DMEM + 2% FBS. Devices were left undisturbed for 1.5 hrs to allow tumor cell attachment. Afterwards, media in all ports was aspirated, and one port for the channel containing the tumor cells was filled with 70 µL of DMEM + 2% FBS while the connecting port was filled with 50 µL of DMEM + 2% FBS. The ports for the opposing media channel were filled with 70 µL and 50 µL of 1:1 DMEM + 2% FBS: EBM-2 + 2% FBS or 1:1 DMEM + 2% FBS: Conditioned Media. For experiments with individual proteins, 50 ng/mL of protein in 1:1 DMEM + 2% FBS: EBM-2 was used to fill the media channel opposite the tumor cells^35-38^. Media was exchanged daily.

### 2.6 Immunofluorescent Staining

Cells encapsulated in microfluidic devices were washed with PBS, fixed using 4% paraformaldehyde, permeabilized using 0.01% Triton X-100 and blocked using 2% abdil. Cells were stained using rat anti-human KI-67 (Cat# 14-5698-82, 1:100) and goat anti-rat Alexa Fluor 555 (Cat# A-21434,1:500), as well as Hoescht 33342 (Cat# 62249, 1:2000). Primary and secondary antibodies were applied during overnight incubation at 4 °C, and devices were washed between antibody incubations using PBS at room temperature. Antibodies and Hoescht were purchased from Thermo Fisher Scientific (Waltham, MA).

### 2.7 Image Acquisition and Analysis

U87-MG migration across the microfluidic device was monitored daily by acquiring fluorescent images of the central gel channel with a DMi8 Yokogawa W1 spinning disk confocal microscope with a Hamamatsu EM-CCD digital camera (Leica Microsystems, Buffalo Grove, IL). Three regions of interest (ROI) were imaged per device. The number of tumor cells that invaded into the central gel channel, along with the distance invaded by each cell, were quantified using the PointPicker plug-in on ImageJ. Per device, the number of invading tumor cells was summed across each ROI, and an average invasion distance was calculated by averaging the distances invaded by all tumor cells imaged per device. The number of invading tumor cells was normalized to unit area.

To compare the number of cells that stain positive for KI-67 between the media channels and gel channels of the microfluidic devices, ImageJ was first used to remove regions containing the gel channel in images containing the media channel as well as the posts in the gel channel images. CellProfiler was then used to identify KI-67+ nuclei in each field of view^39^. The number of KI-67+ nuclei was divided by the number of total nuclei to obtain a percentage. At least two ROI were imaged per media and gel channels per device across three devices.

### 2.8 Temozolomide Dose-Response Assay

Hydrogels containing U87-MG tumor cells were cultured for three days in either 1:1 DMEM + 2% FBS: EBM-2 + 2% FBS or 1:1 DMEM + 2% FBS: Conditioned Media. On Day 3, 500 μM TMZ (Calbiochem, Sigma Aldrich, St. Louis, MO) was added to the media and hydrogels were incubated for an additional 48 hours. 0.5% v/v DMSO was used as a vehicle control. Cell viability for treated and untreated hydrogels was assessed using the MTT assay (Thermo Fisher Scientific, Waltham, MA). The assay was also performed on untreated hydrogels at the beginning of the drug treatment in order to calculate growth rate inhibition (GR_50_) metrics^40^.

### 2.9. Statistics

Statistics were performed using OriginPro (OriginLab, Northampton, MA). Normality of data was determined using the Shapiro-Wilk test, and equality of variance was determined using Levene’s test. For normal data, comparisons between two groups were performed using a t-test, while comparisons between multiple groups were performed using a one-way ANOVA followed by Tukey’s post-hoc when assumptions were met. In the case where data was not normal or groups had unequal variance, comparisons between two groups were performed using a Mann-Whitney test, while comparisons between multiple groups were performed using a Kruskal-Wallis test with Dunn’s post-hoc. Significance was determined as p < 0.05. All quantitative analyses were performed on hydrogels or microfluidic assays set up across at least three independent experiments.

## 3. Results

### 3.1. Conditioned media modulates U87-MG cell number and metabolic activity

We first investigated if conditioned media from three-dimensional endothelial networks would affect the number and metabolic activity of U87-MG cells cultured in GelMA hydrogels for up to a week (**Figure 2**). Endothelial networks were formed via co-cultures of endothelial cells and fibroblasts in 5 wt% GelMA hydrogels (60% DOF) for seven days, with conditioned media collected after this culture period. At Day 3, hydrogels cultured in *CM* had increased numbers of cells compared to those cultured in *EBM2*. Conversely, metabolic activity was decreased in *CM* hydrogels compared to *EBM2* hydrogels. No statistically significant differences in cell number or metabolic activity were observed between experimental groups after extended culture (Day 7).

**Figure 2.**
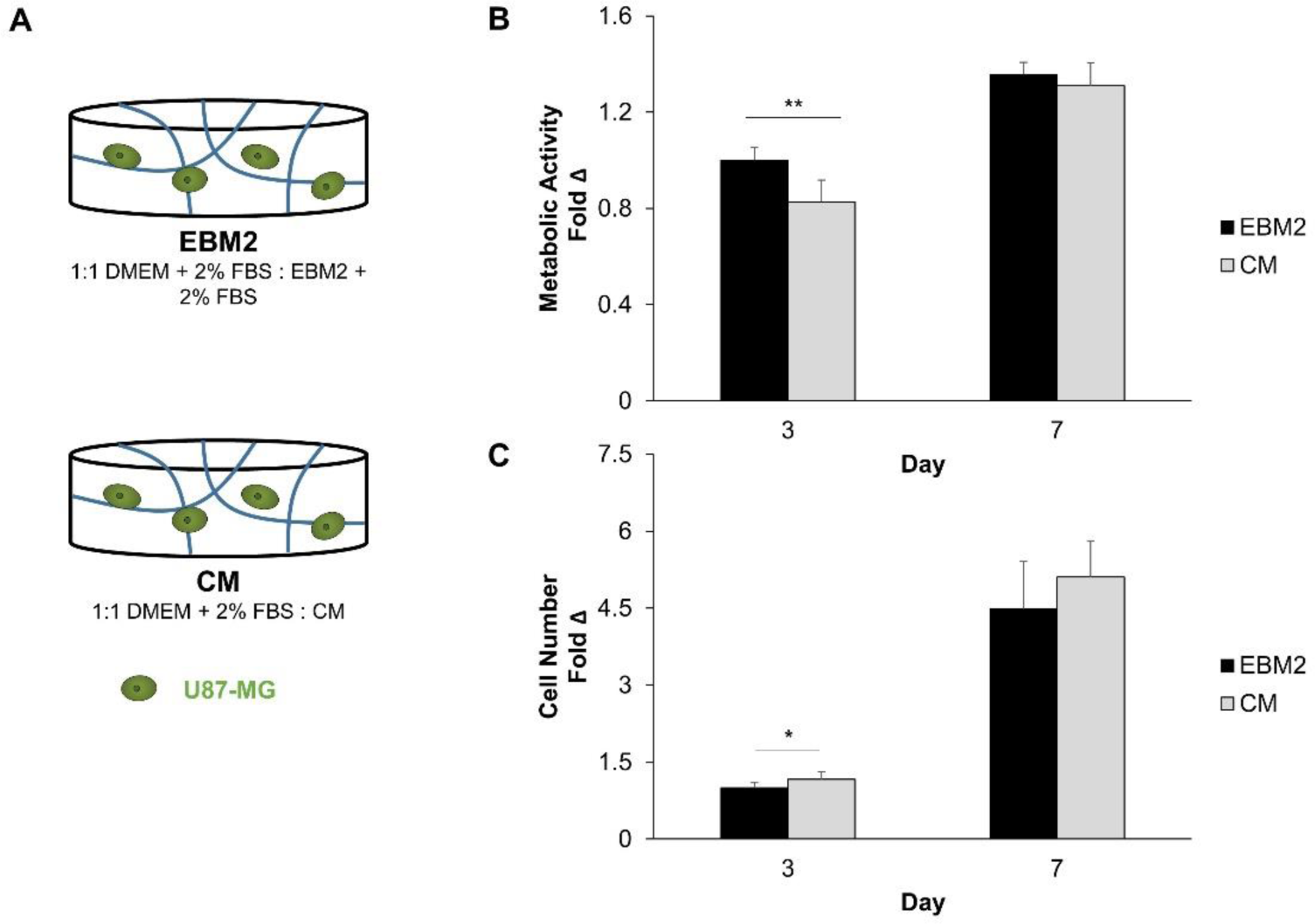
A) Experimental setup to assess the effect of conditioned media on U87-MG metabolic activity (B) and proliferation (C). Metabolic activity (N = 9 hydrogels) and cell number (N = 12 hydrogels at Day 3, N = 9 hydrogels at Day 7) were expressed at each time point as a fold change compared to Day 0. Fold change values were then normalized to the EBM2 group at Day 3. * p< 0.01, ** p<0.001 between groups.

### 3.2. Conditioned media promotes U87-MG migration in a microfluidic invasion assay

We subsequently examined whether the conditioned media contained factors that promoted U87-MG migration. We developed a microfluidic invasion assay using 3D culture chips sold by AIM Biotech (**Figure 3A**). We polymerized GelMA in the central channel of the microfluidic device and then introduced U87-MG cells into one of the side media channels. In the opposing media channel, we introduced either basal (*EBM2*) or conditioned (*CM*) media formulations. We monitored migration of the tumor cells over 72 hours with daily media changes. There were no statistically significant differences in the number of cells remaining in the cell-loading media channel between groups after 72 hours. (**Figure S1**). We found that more cells migrated into the central channel under the influence of *CM*, and that *CM* also increased the average invasion distance of these cells compared to *EBM2* (**Figure 3B**). This suggested that the conditioned media from endothelial networks contained factors promoting U87-MG GBM invasion.

**Figure 3.**
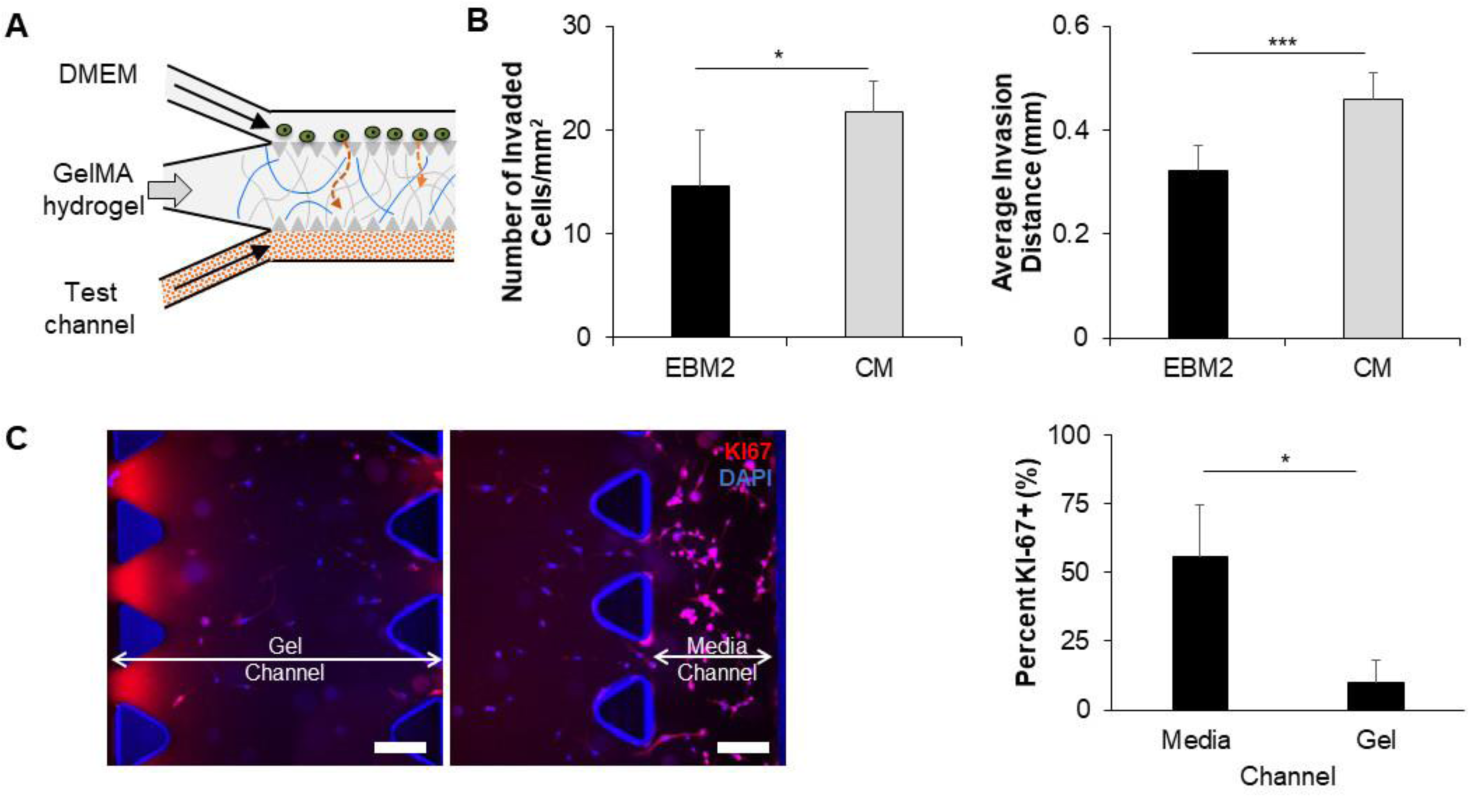
A) Schematic for a microfluidic assay to assess chemotactic cell migration. B) The presence of conditioned media in the test channel leads to increases in the number of cells that migrate into the central gel channel, as well as the average invasion distance of these cells. * p<0.01 between groups, *** p<0.0001 between groups. N = 9 devices. C) Non-migratory cells express KI67 more frequently than migrating cells. Scale bar = 200 µm, * p<0.05 between groups. N = 3 devices.

Additionally, we performed immunofluorescent staining for KI67 within the microfluidic devices to determine whether migrating cells were proliferating. We defined migrating tumor cells as those that were present in the central gel channel, while non-migrating cells were those that remained in the media channel used for seeding. Under the influence of *CM*, tumor cells that did not migrate expressed KI67, while few migrating cells expressed KI67, suggesting that migrating cells were not proliferating (**Figure 3C**). Quantitative comparison of the percent of KI67+ cells between the media and gel channels confirms a significant increase in KI67 expression in cells in the media channel compared to the gel channel. Under the influence of *EBM2*, differences in KI67 expression between the media and gel channels were not statistically significant (**Figure S2**).

### 3.3. Identification of secreted proteins that promote U87-MG migration

We next sought to characterize the composition of the conditioned media in order to identify potential proteins contributing to a chemotactic migratory response. Using a cytokine array, we determined the identities of nine proteins present in the conditioned media based upon the intensities of the corresponding cytokine spots (> 10% of the average intensity of the positive reference spots) (**Figure 4**). The intensities for all proteins except for HGF was significantly higher in *CM* when compared to *EBM2*. We used GlioVis, a data portal containing high-throughput sequencing datasets for brain tumors, to determine whether gene expression of these proteins was differentially regulated in human glioblastoma samples (vs. normal brain) from The Cancer Genome Atlas project (**Table 1**). *IGFBP2, PTX3, SERPINE1, TIMP1*, and *PLAU* were found to be upregulated in glioblastoma samples compared to normal brain samples. In contrast, *HGF* and *THBS1* showed no difference in expression, while *SERPINF1* was downregulated in glioblastoma samples. We additionally performed a literature search to determine potential contributions of each protein to glioblastoma progression (**Table 2**). Several proteins were either found to be over-expressed in perivascular regions of glioblastoma, or implicated in tumor cell proliferation, migration, therapeutic resistance, or cancer stem cell maintenance.

**Table 1.** Comparison of gene expression between GBM tissue specimens from The Cancer Genome Atlas and normal brain tissue specimens using GlioVis

| Gene | Expression in GBM compared to normal brain |
| --- | --- |
| *THBS1* | No difference |
| *SERPINE1* | Upregulated |
| *TIMP1* | Upregulated |
| *SERPINF1* | Downregulated |
| *PLAU* | Upregulated |
| *CXCL8* | Not available |
| *IGFBP2* | Upregulated |
| *PTX3* | Upregulated |
| *HGF* | No difference |

**Table 2.**
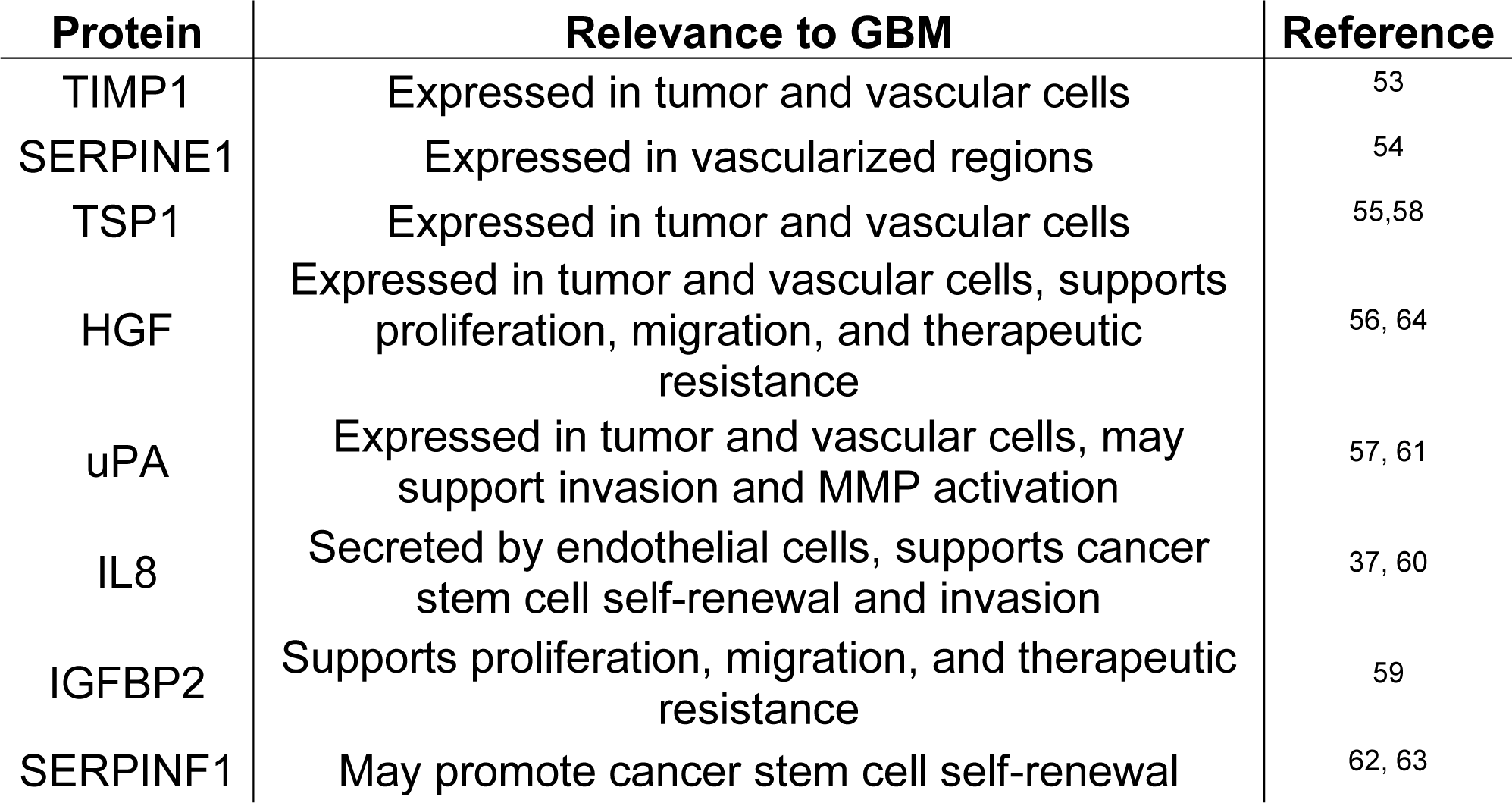
Potential roles of proteins in glioblastoma

| Protein | Relevance to GBM | Reference |
| --- | --- | --- |
| TIMP1 | Expressed in tumor and vascular cells | 53 |
| SERPINE1 | Expressed in vascularized regions | 54 |
| TSP1 | Expressed in tumor and vascular cells | 55,58 |
| HGF | Expressed in tumor and vascular cells, supports proliferation, migration, and therapeutic resistance | 56, 64 |
| uPA | Expressed in tumor and vascular cells, may support invasion and MMP activation | 57, 61 |
| IL8 | Secreted by endothelial cells, supports cancer stem cell self-renewal and invasion | 37, 60 |
| IGFBP2 | Supports proliferation, migration, and therapeutic resistance | 59 |
| SERPINF1 | May promote cancer stem cell self-renewal | 62, 63 |

**Figure 4.**
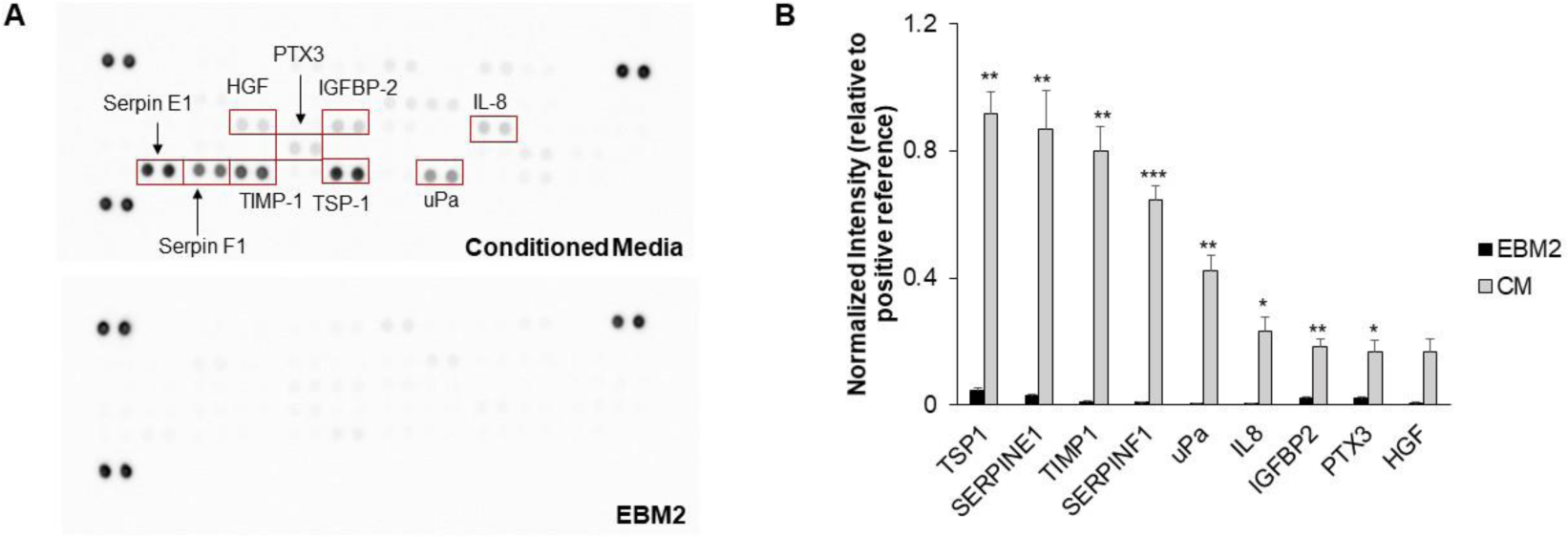
A) Representative images of cytokine array membranes for conditioned media and the basal media control. B) Intensity values for proteins present in conditioned media. The intensities for each protein spot are normalized to the average intensity for the positive reference spots for each membrane. Intensity values for proteins as present in the basal media are also provided for reference. * p< 0.01, ** p<0.001, *** p<0.0001 compared to EBM2. N = 3 - 4 membranes.

Subsequently, we examined the activity of individual proteins identified in the conditioned media to determine which proteins would elicit a chemotactic migratory response from the U87-MG cells. Using the microfluidic invasion assay, we introduced 50 ng/mL of protein in the media channel opposing the channel seeded with tumor cells and monitored invasion for three days. Amongst the proteins whose genes were upregulated in glioblastoma, TIMP1 significantly increased the number of invading cells and the average invasion distance of the tumor cells compared to the negative control of *EBM2* alone (**Figure 5**). We confirmed the presence of TIMP1 in *CM* using an ELISA, which revealed that the TIMP1 was present at a concentration (80 ng/mL) in the same order of magnitude as 50 ng/mL. Proteins whose gene expression were downregulated or not significantly changed between glioblastoma and normal brain specimens did not significantly alter migration compared to the negative control (**Figure S3**).

**Figure 5.**
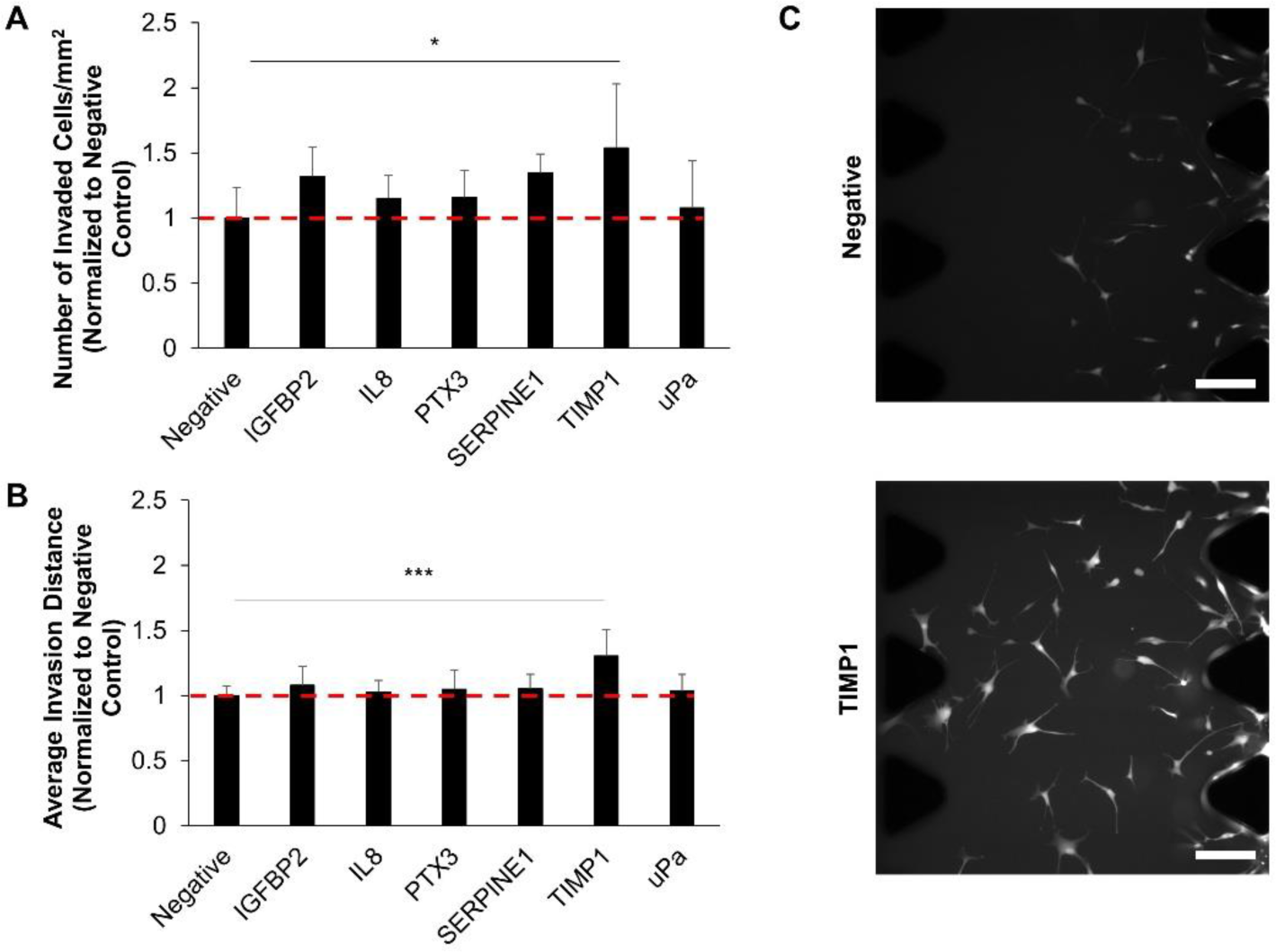
Migration was assessed in response to the presence of each protein loaded into the test channel of the microfluidic assay. The negative control was 1:1 DMEM + 2% FBS:EBM2 + 2% FBS alone. Proteins tested included those whose genes are upregulated in glioblastoma in addition to IL8, which has been shown to induce the migration of primary GBM cells. TIMP1 induced increases in the number of cells (A) and the average invasion distance (B) of cells migrating into the central gel channel. Data is normalized to the Negative group. * p<0.05, *** p<0.001 between Negative and TIMP1 groups. N = 6-9 devices. C) Representative images of cells in the central gel channel. Scale bar = 200 µm.

### 3.4 Conditioned media alters U87-MG response to temozolomide

Finally, we assessed whether U87-MG cells cultured in *CM* displayed a differential response to temozolomide compared to cells cultured in *EBM2* (**Figure 6A**). After culturing hydrogels for three days in either *EBM2* or *CM* media formulations, hydrogels were treated with either 500 µM temozolomide or DMSO for 48 hours before viability was assessed using an MTT assay. To account for the varying growth conditions between *EBM2*- and *CM*-cultured cells, we assessed drug response using growth rate metrics, which normalizes drug response to a single cell division and accounts for the initial conditions of the cultures at the time of treatment. Cells cultured in *CM* had a higher growth rate value compared to those cultured in *EBM2*, suggesting that growth rate was less inhibited by temozolomide in *CM* conditions compared to *EBM2* (**Figure 6B**).

**Figure 6.**
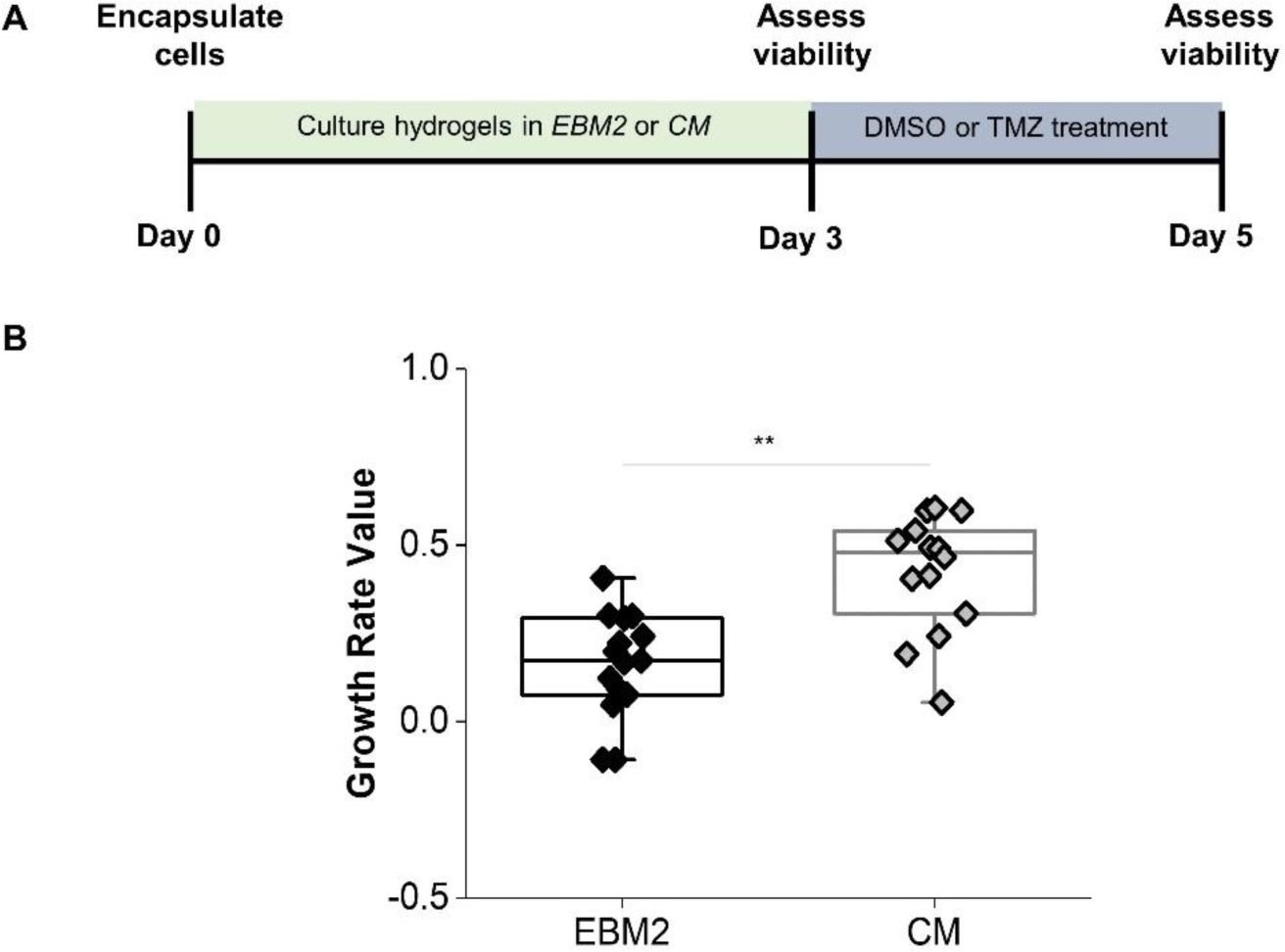
Cells cultured in conditioned media have an increased growth rate value compared to those cultured in basal media after temozolomide treatment. ** p<0.001 between groups, N = 14 - 15 hydrogels.

## 4. Discussion

Glioblastoma tumor cells have been shown to reside in perivascular microenvironments, which facilitates tumor cell migration along vascular tracks, harbors a stem-like subpopulation, and supports differential response to therapy such as radiation^6,16,41^. How the perivascular niche promotes these phenotypes is largely unknown. Studies performed in other cancer types as well as stem cell and regenerative biology reveal a role for angiocrine, or vascular-derived, signals in influencing disease progression, stem cell fate, or regenerative outcomes^42-45^. However, the identities and roles of angiocrine signals in influencing glioblastoma cell behavior is largely unexplored. Vascular cells (e.g. endothelial and mural cells) can communicate with tumor cells via juxtacrine or paracrine signaling, as well as by remodeling or depositing cues in the surrounding extracellular matrix. In this study, we focus on the role of cues secreted by vascular cells in influencing tumor cell proliferation, metabolic activity, invasion, and therapeutic response. Elucidating the effect of paracrine signals on tumor cell behavior enables further investigation into strategies to inhibit the microenvironmental impact on tumor progression. In this manuscript, angiocrine signals are those secreted by both the endothelial cells and fibroblasts in the culture model, in which fibroblasts act as a perivascular mural cells to support network development^25^.

While Transwell assays and direct co-cultures with glioblastoma tumor cells and endothelial cells have been studied, these models often lack the three-dimensionality of the native tissue and fail to recapitulate the formation of structures resembling vasculature. Here, we use GelMA hydrogels to encapsulate both tumor and vascular cell types, and these hydrogels have been shown previously by our lab to modulate tumor cell growth rate as well as promote gene expression, protein secretion, and therapeutic response that favors glioblastoma progression^46-49^. Furthermore, we have shown that GelMA hydrogels support the development of three-dimensional endothelial networks formed from endothelial cells and fibroblasts^32^. HUVECs and NHLFs were chosen because they have been shown to undergo extensive vasculogenesis in various biomaterial systems, and fibroblasts behave similarly to perivascular mural cells in these cultures^23,25,28^. We acknowledge that a limitation of this study is the use of model vascular cell types, which may behave differently from brain-derived sources. Ongoing work is focused on generating vasculature using brain-derived primary vascular cell types in GelMA hydrogels. Our choice of gelatin as the building block for GelMA hydrogel was inspired by the diversity of fibrillar ECM proteins (*e.g*., collagens and fibronectin) tin the perivascular niche^50^. We have shown that GelMA hydrogels can be fabricated such that their elastic modulus is tunable within the 1 – 10 kPa range^31,32^ reported by *Miroshnikova et al*. as the elastic modulus of human glioblastoma specimens via atomic force microscopy^51^.

We identified nine proteins secreted by the endothelial networks self-assembled in GelMA hydrogels. This was based on identifying cytokine spots whose intensities were greater than 10% of the average intensities of the positive reference spots in the cytokine array, because any proteins that were present in EBM-2 had intensities less than 10% (data not shown). In order to determine the relevance of these proteins in GBM biology, we first compared the expression levels of the genes that encode for these proteins between normal brain tissue and GBM specimens from The Cancer Genome Atlas^52^. Most of the identified genes were upregulated in GBM, which might suggest relevance in GBM progression. We additionally searched the literature to determine if these proteins were relevant to GBM. IL8, TIMP1, SERPINE1, HGF, uPa, and TSP1 have been found to be expressed in vascular cells or vascularized regions of human GBM tissue^37,53-58^. IGFBP2 and IL8 have been shown to support GBM proliferation, migration, and to modulate therapeutic response^37,59,60^. uPa may additionally support GBM migration by activating matrix metalloproteinases and by facilitating extracellular matrix degradation^61^. Furthermore, SERPINF1 and IL8 have been implicated in promoting cancer stem cell self-renewal^37,62,63^. These results demonstrate that the secretome of our three-dimensional endothelial networks are relevant to GBM biology, as secreted proteins have either been shown to localize to perivascular regions of GBM tissue or have been implicated in promoting cell behavior that favors tumor progression. Thus, despite the use of model vascular cell types, this platform is relevant to the GBM perivascular niche environment. We further demonstrate that vascular cells are sources for PTX3, IGFBP2, and SERPINF1. These proteins may therefore support various tumor cell behaviors within the perivascular niche, such as invasion, cancer stem cell maintenance, and therapeutic resistance.

We demonstrate that GBM cells exposed to the secretome of the endothelial networks increased in cell number more than tumor cells exposed to control media, as well as exhibited decreased metabolic activity. In a previous study, we used RNA-seq to demonstrate that transcriptomic signatures are altered by interactions between GBM cells and endothelial networks cultured in the same hydrogel^31^. Differential gene analysis revealed that tumor-vascular interactions, which in this case encompassed paracrine, juxtacrine, and reciprocal ECM changes, downregulated genes related to metabolism and cell cycle progression. Results here confirm at a phenotypic level that paracrine tumor-vascular interactions reduce the metabolic activity of tumor cells. Adding to our understanding of GBM-PVN interactions, the observed increase in tumor cell number in response to PVN-conditioned media in this study does not parallel the downregulation of genes related to cell replication observed in our previous study. This may suggest that direct contact between cell types or cell-matrix interactions may have a greater influence overall in dictating proliferation compared to secreted signals. Alternatively, the increase in cell number may reflect a balance between proliferation, survival, and apoptosis. Indeed, upregulated genes uncovered in our prior RNA-seq analysis were also correlated to regulation of cell death and apoptosis. Furthermore, prior literature demonstrates that IL8, IGFBP2, and HGF support GBM proliferation and survival^37,59,64^. Future work will use flow cytometry to elucidate the precise balance between GBM proliferation, survival, and apoptosis governed by angiocrine signaling.

We previously showed that glioblastoma cells reside in proximity to developing endothelial networks in three-dimensional hydrogel models of the brain microenvironment^32^. Here, we chose to specifically investigate the effect of secreted vascular-derived factors on chemotactic migration of tumor cells. While it is known that GBM tumor cells migrate toward vasculature to initiate vessel co-option^65^, the underlying cues and mechanisms that support this directional migratory behavior remain to be elucidated. In particular, the identification of vascular-derived chemokines that support GBM chemotaxis is under-explored^65^. We confirmed that conditioned media enhances the migration of the U87-MG cells. Migrating cells in the central gel channel of the microfluidic device we used to profile invasion were largely non-proliferative by minimal KI-67 expression, suggesting increased presence of cells in the central gel channel was due to migration from the cell seeding channel and not division of invading cells. This behavior is also indicative of the go-or-grow phenomena observed from brain tumors; this hypothesis suggests that migration and proliferation are mutually exclusive events^66,67^. Prior work from our lab has observed the go-or-grow phenomena under hypoxic conditions, and *Farin et al*. demonstrate that tumor cells invading along a blood vessel will momentarily cease migration in order to divide^7,68^. While other studies have shown that GBM tumor cells will migrate toward endothelial cells, the specific proteins that promote chemotactic migration have not yet been identified^35,69,70^. Based upon prior literature that established migration assays in microfluidic devices or utilized Transwell assays to investigate GBM response to similar proteins^35-38^, we investigated U87-MG migration in response to protein gradients established by 50 ng/mL of each individual protein identified in the conditioned media. Intriguingly, we identified TIMP1 as a potent promoter of U87-MG migration. While TIMP1 is traditionally known as an inhibitor of matrix metalloproteinases, it may also function in matrix metalloproteinase-independent roles as a cytokine by binding to cell surface receptors to affect behaviors such as proliferation, apoptosis, and differentiation^71^. Exogenous TIMP1 has been shown to promote the migration of cancer associated fibroblasts and neural stem cells through CD63 and integrin signaling^72,73^. While prior literature has focused on the effect of directly treating GBM cells with TIMP1 on resulting invasion, our results reveal a role for vascular-derived TIMP1 as a chemotactic chemokine that supports GBM tumor cell migration towards vasculature. Future efforts will investigate whether inhibition of TIMP1 or CD63 reduces the association between endothelial networks and glioblastoma cells in three-dimensional hydrogel co-cultures, as well as determining whether additional GBM cell sources respond to TIMP1 as a chemotactic agent. Additionally, while we screened for the effects of individual soluble factors in modulating U87-MG migration, we are also interested in investigating combinations of soluble factors to determine synergistic effects on migration.

Finally, we demonstrate that paracrine signaling from endothelial networks tempers U87-MG response to temozolomide, the current standard-of-care chemotherapy for glioblastoma. The perivascular niche has been implicated in blunting the effects of therapy on GBM tumor cells^16^; studies have primarily focused on modulating the effect of radiation, but resistance to temozolomide has also been observed^74^. In support of our results, IGFBP2 has been shown to promote temozolomide resistance through integrin β1-ERK signaling^59^. Our findings that angiocrine signaling supports TMZ chemoresistance in glioblastoma are consistent with previous efforts from our group that showed that direct co-culture of GBM cells within endothelial cell networks in GelMA hydrogels are less responsive to temozolomide compared to GBM cells alone. That work also suggested that tumor-vascular interactions modulated *MGMT* and mismatch repair gene expression profiles, which are DNA repair mechanisms responsible for temozolomide resistance^31^. As a result, ongoing efforts are examining if the other proteins identified in this study are individual mediators of temozolomide resistance in order to identify underlying signaling mechanisms that lead to this response. Potential mechanisms include modulation of MGMT and mismatch repair functionality, as well as integrin-mediated resistance^13^.

## 5. Conclusions

The role of soluble angiocrine-derived signals in glioblastoma progression remains to be fully explored and may contribute to behaviors such as invasion and therapeutic resistance as observed in the perivascular niche environment. In this paper, we demonstrated the adaptation of tissue engineering platforms to demonstrate that endothelial networks self-assembled in gelatin hydrogels secrete proteins that are relevant to glioblastoma biology. These secreted factors supported GBM cell proliferation, migration, and therapeutic resistance to temozolomide. Intriguingly, we identified a novel role for TIMP1 in mediating directional migration of U87-MG cells, which indicates a potential role in initiating vessel co-option in glioblastoma. Overall, these results provide insight into the role of paracrine tumor-vascular signaling in glioblastoma progression and highlight the relevance of biomaterial-based vascular models in investigating glioblastoma biology.

## Acknowledgements

The authors gratefully acknowledge Professor Roger Kamm, Dr. Giovanni S. Offeddu, and Dr. Kristina Haase at MIT for technical discussions that aided experiment execution. This work was supported by the National Cancer Institute of the National Institutes of Health (R01 CA197488) and the National Science Foundation Graduate Research Fellowship (DGE 1144245 to MTN). The content is solely the responsibility of the authors and does not necessarily represent the official views of the NIH. The authors also acknowledge additional funding provided by the Department of Chemical and Biomolecular Engineering and the Carl R. Woese Institute for Genomic Biology at the University of Illinois at Urbana-Champaign.

