## Supplemental Figures 1-3 for "Multidimensional hydrogel models reveal endothelial network angiocrine signals increase glioblastoma cell number, invasion, and temozolomide resistance"

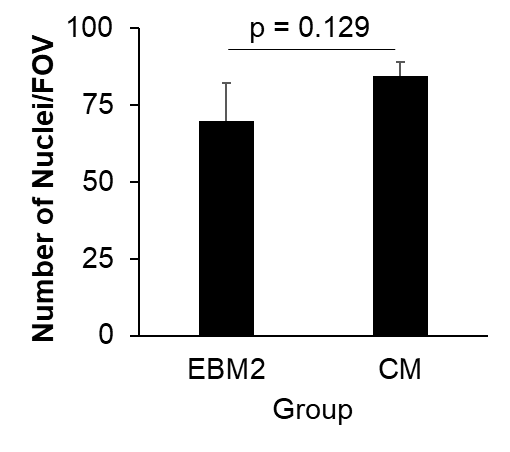

**Figure S1**. Cell loading consistency between *EBM2* and *CM* groups was assessed by imaging the media channel containing cells after 72 hours and counting the number of nuclei per field of view (FOV). N = 3 devices.

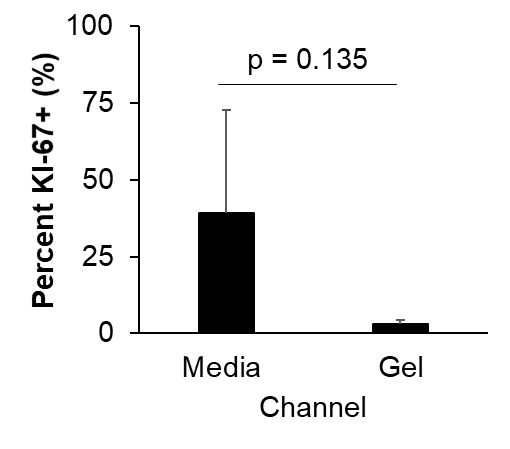

**Figure S2.** When 1:1 DMEM + 2% FBS : EBM-2 + 2% FBS was used in the test channel of the invasion assay, the difference in the percent of KI67-positive cells between the media and gel channels was not statistically significant. N = 3 devices.

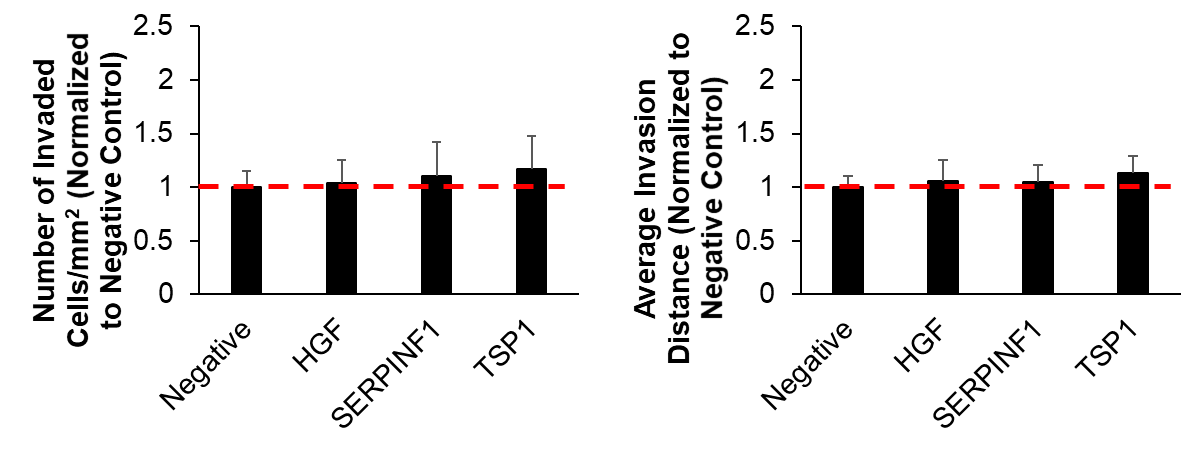

**Figure S3.** Migration was assessed in response to the presence of each protein loaded into the test channel of the microfluidic assay. The negative control was 1:1 DMEM + 2% FBS : EBM2 + 2% FBS alone. Proteins were those downregulated or not differentially regulated in glioblastoma compared to normal brain tissue, as determined using GlioVis to analyze data from The Cancer Genome Atlas. Data is normalized to the Negative group.
